# InstaNovo-P: A de novo peptide sequencing model for phosphoproteomics

**DOI:** 10.1101/2025.05.14.654049

**Authors:** Jesper Lauridsen, Pathmanaban Ramasamy, Rachel Catzel, Vahap Canbay, Amandla Mabona, Kevin Eloff, Paul Fullwood, Jennifer Ferguson, Annekatrine Kirketerp-Møller, Ida Sofie Goldschmidt, Tine Claeys, Sam van Puyenbroeck, Nicolas Lopez Carranza, Erwin M. Schoof, Lennart Martens, Jeroen Van Goey, Chiara Francavilla, Timothy Patrick Jenkins, Konstantinos Kalogeropoulos

## Abstract

Phosphorylation, a crucial post-translational modification (PTM), plays a central role in cellular signaling and disease mechanisms. Mass spectrometry-based phosphoproteomics is widely used for system-wide characterization of phosphorylation events. However, traditional methods struggle with accurate phosphorylated site localization, complex search spaces, and detecting sequences outside the reference database. Advances in *de novo* peptide sequencing offer opportunities to address these limitations, but have yet to become integrated and adapted for phosphoproteomics datasets. Here, we present InstaNovo-P, a phosphorylation specific version of our transformer-based InstaNovo model, fine-tuned on extensive phosphoproteomics datasets. InstaNovo-P significantly surpasses existing methods in phosphorylated peptide detection and phosphorylated site localization accuracy across multiple datasets, including complex experimental scenarios. Our model robustly identifies peptides with single and multiple phosphorylated sites, effectively localizing phosphorylation events on serine, threonine, and tyrosine residues. We experimentally validate our model predictions by studying FGFR2 signaling, further demonstrating that InstaNovo-P uncovers phosphorylated sites previously missed by traditional database searches. These predictions align with critical biological processes, confirming the model’s capacity to yield valuable biological insights. InstaNovo-P adds value to phosphoproteomics experiments by effectively identifying biologically relevant phosphorylation events without prior information, providing a powerful analytical tool for the dissection of signaling pathways.

## Introduction

Phosphorylation is a widespread protein post-translational modification (PTM) and the most abundant PTM in eukaryotes^1–3^. It plays crucial roles in enzyme activation, signal transduction, and transcription regulation, with aberrant activity linked to various pathological states, including cancer^4–6^. Consequently, detection of phosphorylation events is imperative for dissecting cellular pathways and describing disease states. Kinases, the enzymes responsible for phosphorylation, are significant drug targets and have found considerable success in therapeutic intervention^7,8^.

Mass spectrometry has become the *de facto* approach for studying protein phosphorylation across different cell types and tissues under various conditions^9–13^. Phosphoproteomics, the field dedicated to studying phosphorylated proteins, has seen major advancements in sample preparation, instrumentation, and data analysis for phosphorylated peptide detection over the past decades^14–20^. The traditional approach for identifying phosphorylated peptides involves database searches based on the mass shift induced by phosphorylation in the sequence search space. However, this method has inherent limitations in modification localization ambiguity, and significantly expands the search space, search time and costs^21–23^. Additionally, target-decoy methods cannot detect non-canonical protein sequences or protein isoforms that are not included in the database search, restricting analysis to only known reference proteomes.

Recent advances in *de novo* peptide sequencing using machine learning approaches have shown impressive performance in peptide detection, including peptides with modifications such as N-terminal acetylation and methionine oxidation^24–26^. These methods have the potential to address the shortcomings of database searches, provide complementary information, and be integrated into hybrid approaches to maximize insights into phosphorylation in investigated proteomes. Despite the abundance of publicly available PTM proteomics datasets, *de novo* peptide sequencing approaches have not been extensively adapted for PTM identification.

Previous transformer-based methods for *de novo* peptide sequencing, including InstaNovo^24^, predominantly utilize auto-regressive architectures, predicting amino acids sequentially. Recently, Xia et al. developed AdaNovo^27^, incorporating conditional mutual information between spectra and peptides, successfully identifying amino acids carrying PTMs on benchmark datasets, however without including phosphorylation. PrimeNovo^28^, a model using non-autoregressive prediction, is currently the best model that supports detection of phosphorylated peptides.

Here, we introduce the first application of a *de novo* peptide sequencing model that is trained on an extensive number of phosphoproteomic datasets and is exclusively adapted for phosphorylated peptide detection. We reprocess numerous phosphoproteomics experiments available in the PRIDE database^29^ to make a fine-tuned, phosphorylation-specific version of InstaNovo, achieving substantial improvements in phosphorylated peptide identification accuracy. We validate the performance of our fine-tuned model across multiple datasets, and confirm its predictions with targeted proteomics. Lastly, we provide for the first time experimental validation of phoshorylated peptide predictions that went undetected by conventional database searches.

## Results

### InstaNovo-P is a phosphorylation-specific de novo sequencing model

We utilized our previously developed *de novo* sequencing model, InstaNovo^24^, and adapted it to the task of phosphorylated peptide detection. InstaNovo is an autoregressive transformer model with an encoder-decoder architecture, trained on the ProteomeTools dataset^30^. During inference, InstaNovo interprets an MS/MS spectrum by translating it to the peptide sequence that produced the spectrum. To fine-tune InstaNovo for phosphorylated peptide prediction (i.e., to further train and adapt our model for the specific task of phosphorylated peptide sequencing), we curated a set of 29 reprocessed phosphoproteomics projects from Scop3P^31^ (see Supplementary Table 1 for complete list of projects). After filtering for high localization probability (*>*0.80), our training dataset consisted of 2,760,939 PSMs mapping to 74,686 unique peptides. Three projects made up more than half of the PSMs with PXD000612 alone containing 837,540 or 31% of the PSMs (Figure 1A). When accounting for unique peptides per project, the analysed dataset was also dominated by a few large projects, although the distribution is slightly more even (Figure 1B). The distribution of phosphorylated amino acids is dominated by serine (S), making up 84.5%, while threonine (T) and tyrosine (Y) make up 12.0% and 3.4%, respectively (Figure 1C). To minimize data leakage, the dataset was split using GraphPart^32^ into a train-valid-test split of 70-10-20%.

**Figure 1.**
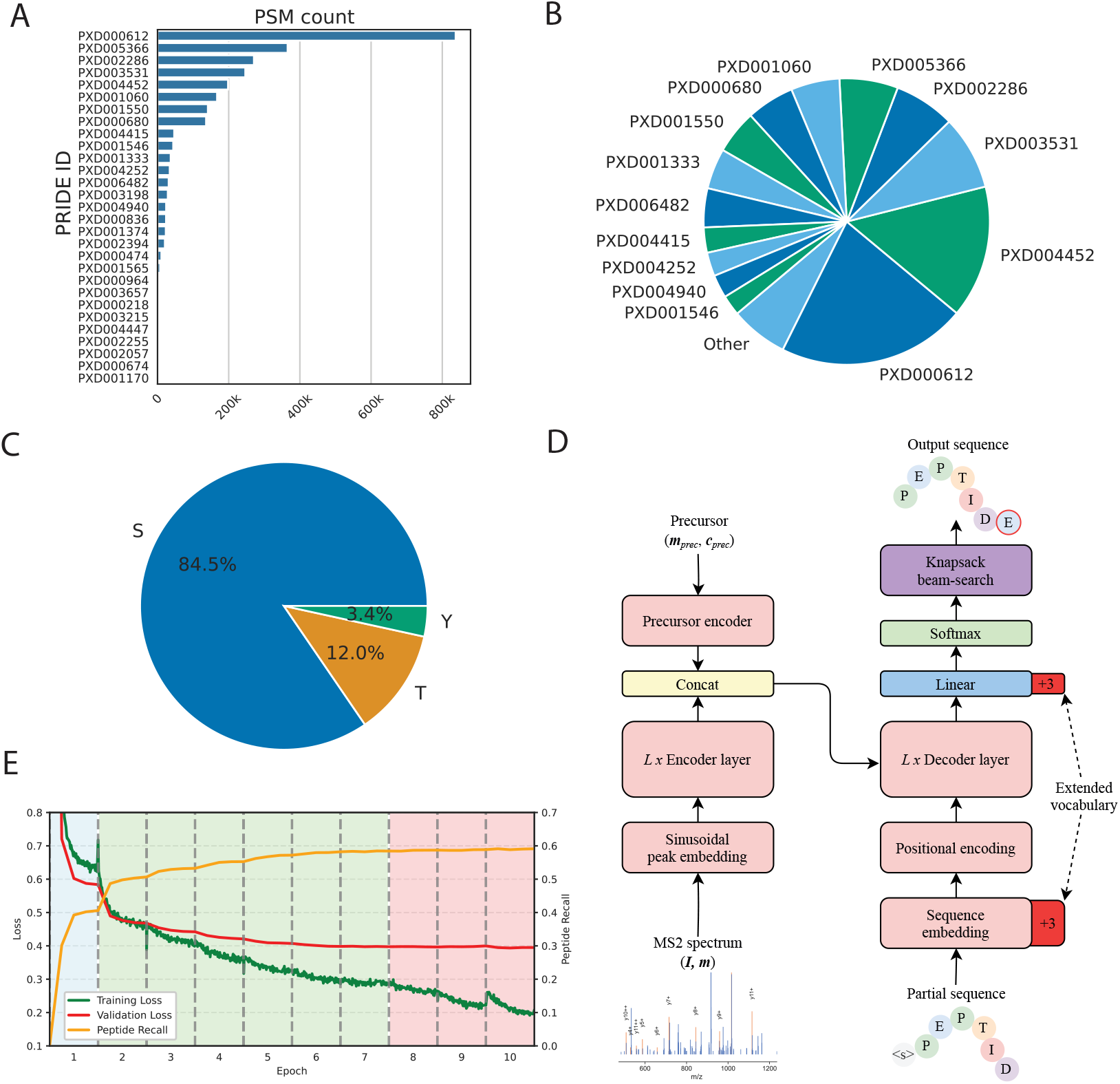
Overview of training data and InstaNovo-P model. A) PSMs per PRIDE project reprocessed and used to train the data. B) Unique phosphorylated peptide sequences used to train the model per project. C) Percent of phosphorylated residue types in the combined train-valid-test set. D) InstaNovo-P architecture. The sequence embedding and linear prediction layer in the decoder head have been adapted to accommodate the prediction of log probabilities for phosphorylated tyrosine, serine and threonine tokens, in addition to the standard vocabulary and modifications of the base InstaNovo model. E) Training loss, validation loss, and peptide recall during fine-tuning with gradual unfreezing. The blue region had only the head and embedding layers unfrozen, the middle greed region was gradually unfrozen in the encoder layers, the final red region was gradually unfrozen in the decoder layers.

We used the InstaNovo model architecture with two modifications. The sequence embedding layer and the linear head of the model had to be extended to accommodate three extra tokens for the possible phosphorylated residues of serine, threonine, and tyrosine (Figure 1D). To fine-tune, we utilized gradual unfreezing^33^ of the model layers starting with the head and sequence embedding for 1 epoch. Then, instead of unfreezing the decoder, we gradually unfroze the encoder layers. Finally, the decoder layers were also gradually unfrozen, reducing the learning rate with increasing number of training steps (Supplementary Figure S1A). Interestingly, the model reached good validation performance after only fine-tuning the head, sequence embedding and encoder, but the decoder is also fine-tuned for the last performance gain (Figure 1E, see Methods for further details).

To assess the model performance on phosphorylated peptides, we used 3 evaluation datasets: the test set made from the split of the curated projects, here referred to as “Test”, a subset of project PXD009449^34^, here referred to as “21PTM”, and a dataset from an in-house experiment using T47D breast cancer cell line expressing or not Fibroblast Growth Factor Receptor 2 (FGFR2) upon growth factor treatment, here referred to as “FGFR2”. The three evaluation datasets contained different distributions of non-phosphorylated and phosphorylated peptides as well as different residue types (Supplementary Figure S1B and S1C). In addition to the evaluation on phosphorylated datasets, we also evaluate the model on 2 datasets of unmodified peptides: The original test set used by the base model, here referred to as “AC-PT”, and the unmodified version of 21PTM, here referred to as “21PTM-unmod”.

In summary, we extended the InstaNovo transformer by adding three phosphorylation-specific tokens and fine-tuned on a 2.76 million PSM phosphoproteomics corpus, achieving strong validation performance using an encoder-first gradual unfreezing schedule.

### InstaNovo-P assigns phosphorylation events with high accuracy and precision

We compared the performance of our model with the base InstaNovo model (version 0.1.4) on unmodified peptides from the AC-PT dataset (ProteomeTools Parts I-III)^24^ and the 21PTM dataset^34^. InstaNovo-P increases peptide recall from 64.0% to 68.1% and from 70.0% to 71.4% for the AC-PT test set and 21PTM dataset (taking into account only unmodified peptides), respectively, compared to the base InstaNovo model (Figure 2A). This surprising observation indicates that the fine-tuning effort did not constrain the model to the detection of phosphorylated peptides only, but also improved the general performance. In other words, the model did not experience catastrophic forgetting even without mixing training data from target and source domain (i.e., unmodified vs modified peptides).

**Figure 2.**
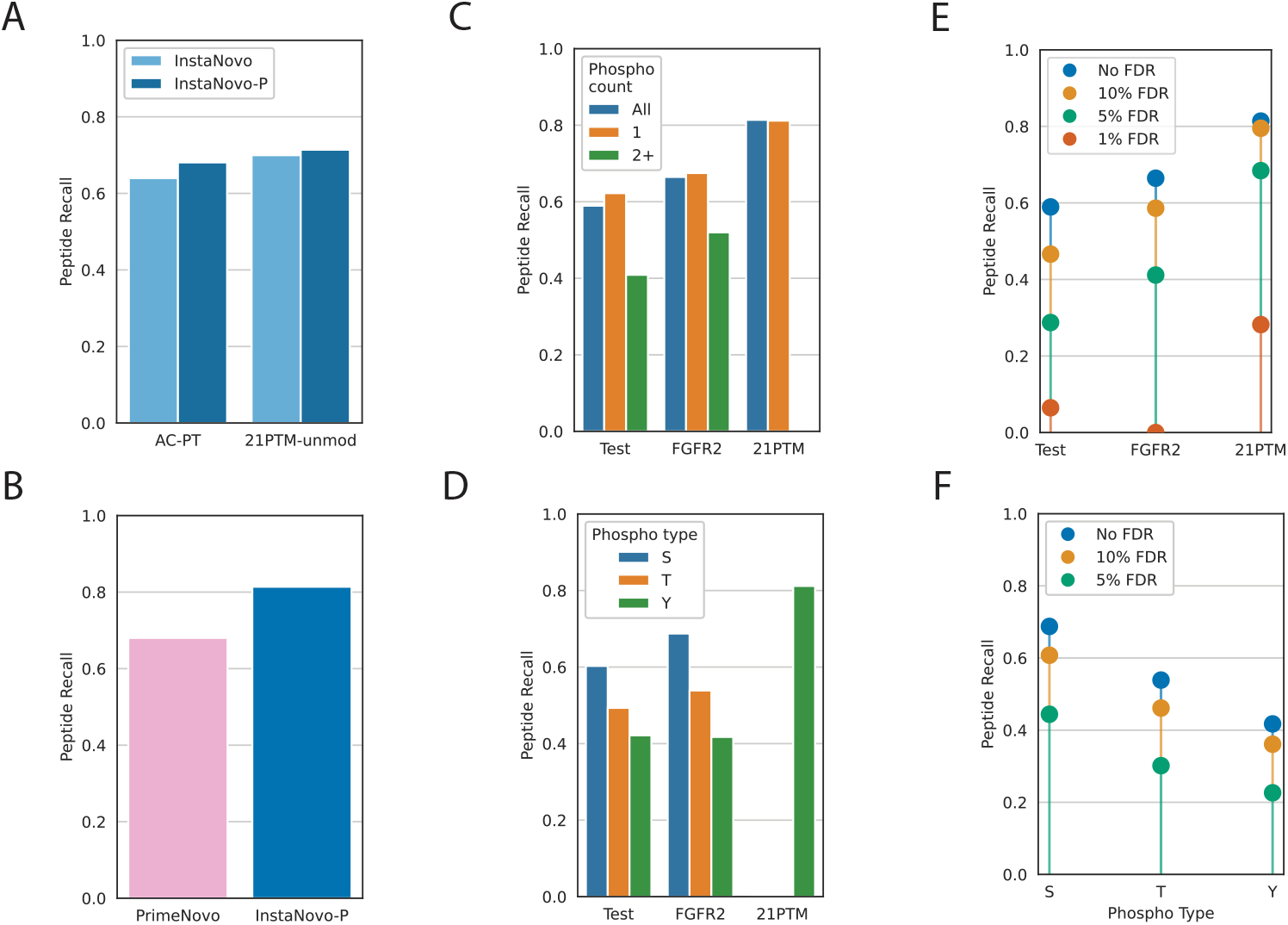
Benchmarking and evaluation of InstaNovo-P predictions. A) Comparison of Instanovo and InstaNovo-P peptide recall on the AC-PT (Proteome Tools I-III) and 21PTM dataset. B) Comparison of PrimeNovo and InstaNovo-P peptide recall on the 21PTM dataset. C) Peptide recall for each test dataset for all peptides, and peptides with different number of phosphorylated sites per peptide. D) Peptide recall for each test dataset split by type of phosphorylated residue. E) Peptide recall in each dataset for different FDR thresholds as calculated by database grounding. F) Peptide recall for different FDR thresholds, split by phosphorylated residue type in the FGFR2 dataset.

We then evaluated the performance on phosphorylated peptides of the 21PTM dataset. InstaNovo-P achieved a peptide recall of 81.4% (Supplementary Table 2). In comparison, PrimeNovo^28^ reported a peptide recall of 68% (Figure 2B). This increase of 13.4% compared to PrimeNovo indicates an excellent generalization to unseen phosphorylated peptides. Moreover, PrimeNovo was fine-tuned specifically on 21PTM, whereas InstaNovo-P evaluated 21PTM entirely as a holdout dataset.

Next, we evaluated the performance of our model in relation to the number of phosphorylated sites present on phosphorylated peptides. Our evaluation datasets contain unmodified peptides, as well as peptides with 1, 2, or more phosphorylation sites (Supplementary Figure S2A). As expected, we see decreasing peptide recall with the increased number of phosphorylated sites in each peptide (Figure 2C).

Unsurprisingly, we also observed a correlation of peptide recall with the composition of our training dataset. The highest recall was seen in serine phosphorylation events (Supplementary Table 3), which were also the most abundant events in our training dataset (Figure 2D). The highest recall was seen for the 21PTM dataset, which contained only phosphorylated tyrosines. We speculate that the reason is due to high intensity signal, the nature of the synthetic peptides, and lower sample complexity. Limiting and estimating the number of false positives in proteomics searches is imperative for utility in biological studies and downstream validations. We therefore evaluated the model confidence threshold needed for different values of false discovery rate (FDR, Supplementary Figure S2B). The confidence range distribution of predictions for all evaluation datasets was similar to the base model, and indicates that the model can inherently distinguish between correct and incorrect assignments (Supplementary Figure S3). We observed highly diminishing peptide recall at 1% FDR (Figure 2E). Thus the best threshold that balanced recall and by extension identification performance with a satisfactory FDR was 5%. We also observed the same trend based on residue phosphorylation type. Peptide recall on the FGFR2 dataset by type of phosphorylated residue at 5% FDR was 44.4%, 30.2% and 22.6% for S, T and Y, respectively (Figure 2F).

Altogether, these data indicate that InstaNovo-P generalizes well to phosphorylated peptides without catastrophic forgetting, while the predicted confidence scores effectively discriminate correct assignments, with a 5% FDR threshold offering the optimal trade-off between recall and reliability.

### InstaNovo-P performance is dependent on peptide properties

We next evaluated the precision and recall of peptide predictions on the three different test datasets. As previously observed for our base InstaNovo model, we saw highly variable performance depending on the dataset evaluated (Figure 3A and B). InstaNovo-P achieved the best performance on 21PTM, which was composed of synthetic peptides. We recorded lower recall in our test and FGFR2 dataset, which originated from other projects with endogenous peptides. We noted that performance was dependent on to peptide length (Figure 3C and D). InstaNovo-P exhibited lower peptide recall with increasing peptide length, which affected the dataset total recall. Even though the overall recall for 21PTM was higher than that of the test set and FGFR2 dataset, we found that the recall for shorter peptides (under 15 amino acids) was almost identical across datasets (Figure 3D). The FGFR2 dataset included both a phospho-enriched and an unmodified version of several peptides. In addition to those two, we also took the phospho-enriched FGFR2 and “de-modified” it (changing the phosphorylated sites to their non-modified version). A prediction would usually contain a phosphorylated site, but by de-modifying it, we could infer the added difficulty of phosphorylation localization alone. We found that InstaNovo-P performed well across all three types, however phosphorylated peptide identification was more difficult than for unmodified peptides (Figure 3E). For longer peptides, the recall performance was noticeably higher for the FGFR2 dataset when compared to the two other evaluation datasets, with recall also showing dependence on the number of phosphorylated sites for each peptide (Figure 3F) .

**Figure 3.**
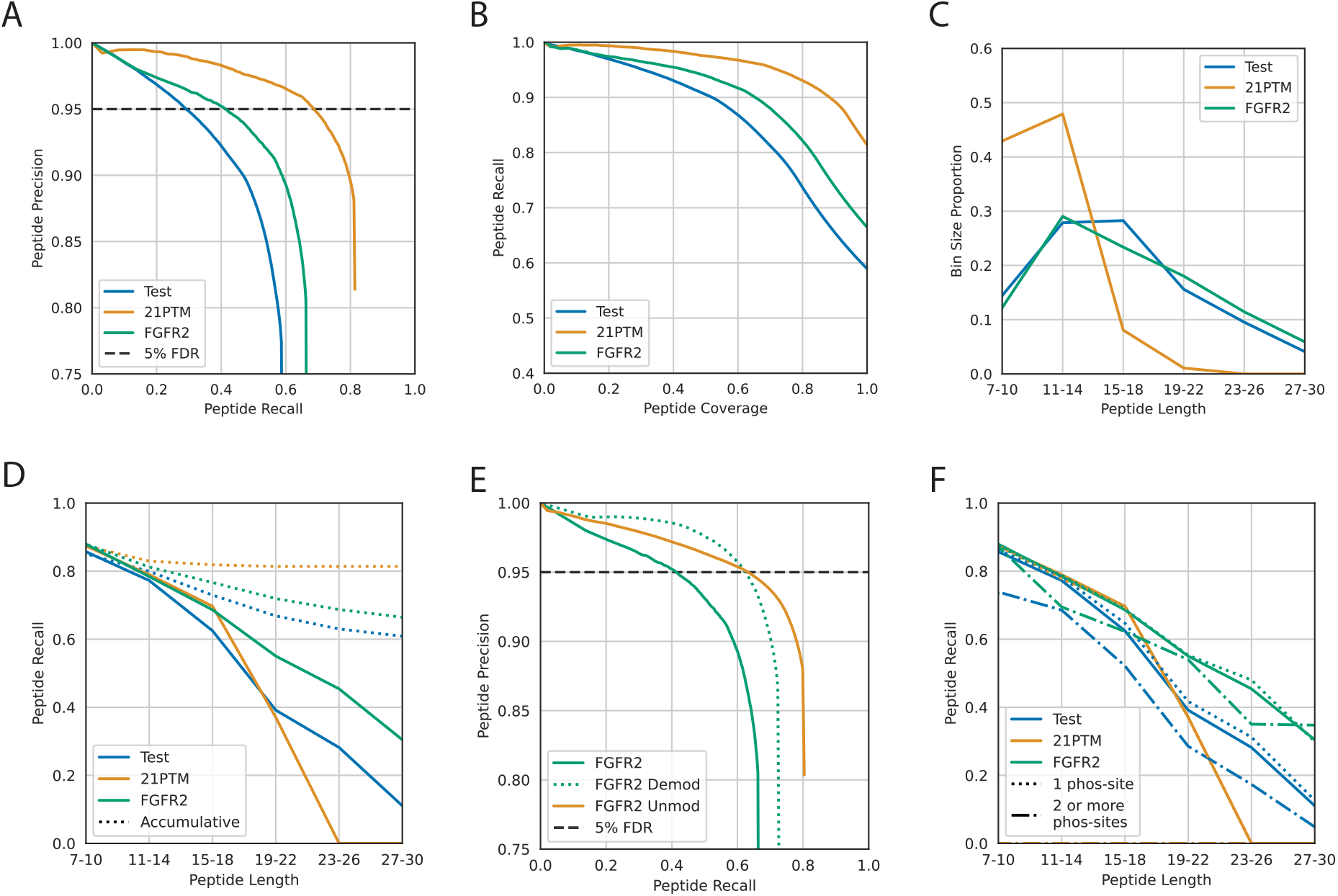
Prediction accuracy is related to peptide properties. A) Precision-recall curves for each test dataset. Horizontal line indicates 5% FDR threshold as calculated by database grounding. B) Recall-coverage curve for each of the test datasets. C) Composition of test datasets as assessed by peptide length. D) Recall per bin for each test dataset. E) Precision recall curve for the FGFR2 dataset including Demod, which has all modifications in both predictions and targets removed, and Unmod, which only contains unmodified peptides. Horizontal line indicates 5% FDR threshold. F) Peptide recall for each test dataset, with separate curves for peptides that have one and two or more phosphorylated sites.

In sum, InstaNovo-P performance varied markedly by dataset and peptide characteristics, while recall also trended downward as the number of phosphorylation sites per peptide increased. Our results suggest that our recall differences between the three evaluated datasets were predominantly due to differences in peptide attributes.

### InstaNovo-P is a robust tool for proteome-wide identification of phosphorylation events

We used our FGFR2 dataset to map our model’s predictions to the proteome, providing another validation for our predictions, given the very low likelihood of an incorrect prediction mapping to the proteome. We also compared our results to database search results using MaxQuant. We assessed our false discovery rate using the database search identifications in the labeled space (scans in common with database results) or the full search space (all scans in the dataset). We noticed that InstaNovo-P predicted substantially more phosphorylated peptides at 5% FDR when evaluated in the full search space rather than the labeled space alone, and that the gains are similar across different perturbations and time points in our experiments. Overall, InstaNovo-P detected 97.1% more peptides (Figure 4A). On average, InstaNovo-P detected 3, 068(*±* 495) peptides, which was expanded to 5, 436(*±* 770) peptides when expanding our search to the full search space, i.e., also scans that the database did not identify (Figure 4B). We mapped these peptides to 21.6% more proteins in the proteome compared to the database search in the same dataset (4B). The identified peptides mapped to 1, 331(*±*209) proteins on average in the labeled space, which increased to 1, 544(*±* 239) proteins in the full search space (Figure 4D).

**Figure 4.**
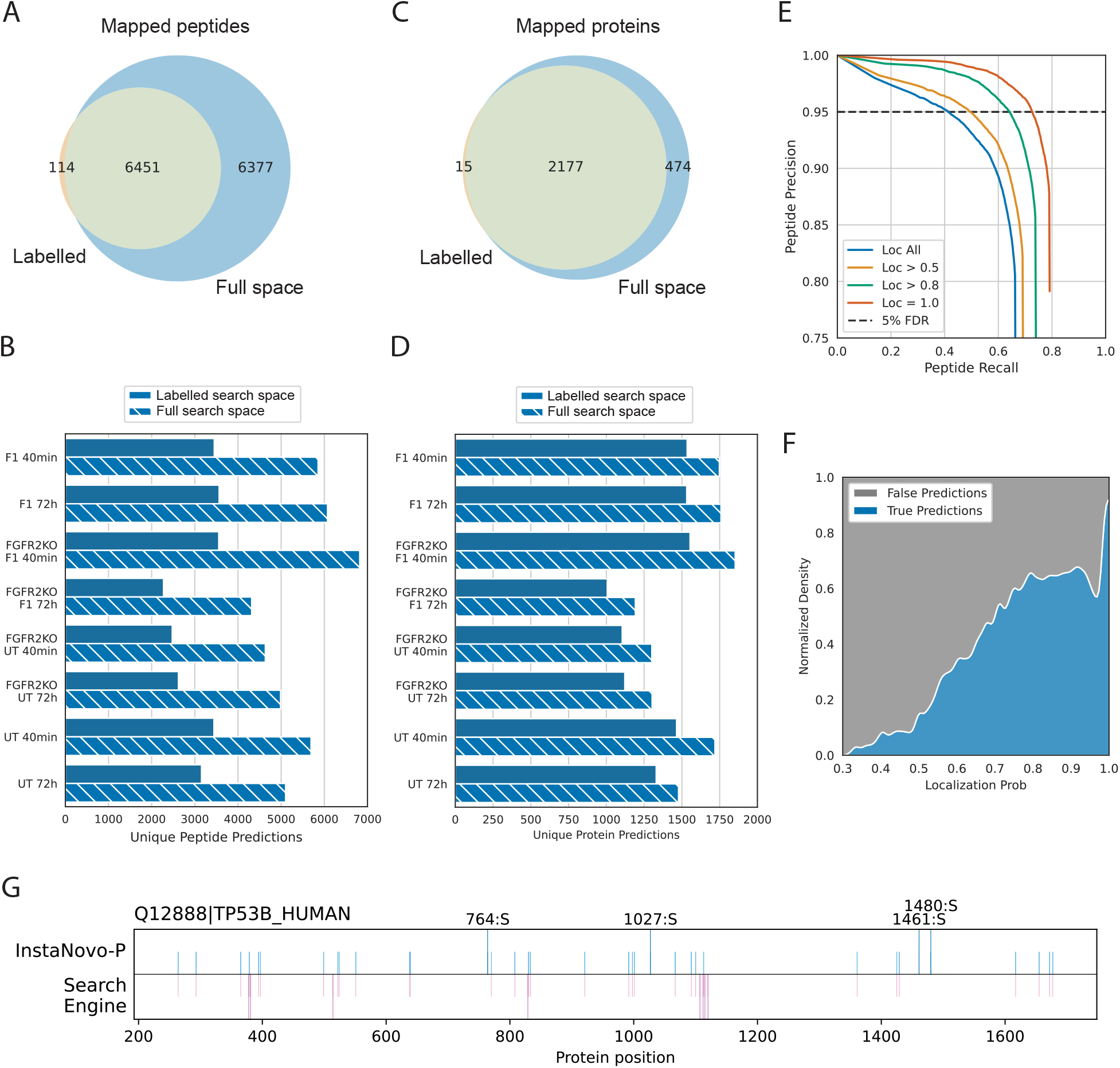
InstaNovo-P expands identifications and localization certainty. A) Venn diagram of the number of phosphorylated peptides in the database search space (labeled), and predicted phosphorylated peptides from InstaNovo-P in full search space of the FGFR2 dataset. B) Proteins detected with database search and InstaNovo-P at an estimated 5% FDR threshold in labeled and full search space. C) Peptides in database search and InstaNovo-P with 5% FDR for each condition in the experiment. D) Proteins detected with database search and InstaNovo-P with 5% FDR for each condition in the experiment. E) Localization precision - peptide recall curve for different localization certainty thresholds according to database search. Horizontal line indicates 5% FDR as calculated by database grounding. F) Normalized density of true positive (True) and false positive (False) predictions, as ranked by localization certainty according to the database search. G) Mapping of phosphorylated sites detected with InstaNovo-P (top) and database search (bottom) in protein TP53 for all experimental conditions. Phosphorylated sites that are unique to either method are extended.

The localisation accuracy of InstaNovo-P, as assessed by recall, indicated that the model is adept at recognizing and predicting phosphorylation events for peptides containing multiple potential phosphorylation sites. The recall observed, as well as the recall at an estimated 5% FDR, positively correlated with database localization certainty (Figure 4E). However, peptides with low localization certainty included in the database search results might contain incorrect localization certainty that negatively affects peptide recall. This is substantiated by the trend of database localization probability as a function of correct and incorrect assignments by InstaNovo-P in the same dataset (Figure 4F).

We then turned our attention to a number of specific phosphorylation sites in proteins that are critical components of the cell cycle. We first investigated phosphorylated peptide predictions for sites in the protein TP53, an important cell cycle checkpoint^35^. InstaNovo-P predicted 35 phosphorylated sites in total, of which 4 contain phosphorylation sites in TP53 that went undetected in the database search space (Figure 4G). InstaNovo-P predicted 31 phosphorylation sites in common with the database search, while the search predicted 8 sites that InstaNovo-P did not detect.

We also evaluated predicted peptides and phosphorylation sites not detected by database search in lamin A/C (LMNA), a protein that is known to be phosphorylated during mitosis and is thought to be involved in chromatin stability and regulation^36^. In the full search space, InstaNovo-P predicted 6 more peptides containing 2 more unique phosphorylated sites that are not present in the labeled space (Supplementary Figure S4).

Collectively, proteome-wide mapping in the FGFR2 dataset shows that InstaNovo-P not only nearly doubles identified phosphorylated peptides at 5% FDR by extending our search to the full search space, but also retains high localization accuracy that scales with database search localization certainty. Importantly, it uncovers novel phosphorylation sites, demonstrating potential in revealing additional phosphorylation sites beyond database searches.

### InstaNovo-P predicts phosphorylation sites that contribute meaningful biological insights

To evaluate potential novel insights provided by our model we used a subset of the FGFR2 dataset and compared T47D cells, either wild type (WT) or CRISPR/CAS9-depleted of FGFR2 (KO), upon treatment with the specific FGFR2 ligand FGF7 (F7)^37^ for different timepoints (40 min or 72 hours). Each of these conditions contained only phospho-enriched samples, eluted in 2 separate fractions per column. We mapped InstaNovo-P predictions from these experimental files in the human proteome, and assessed their presence in the labeled space. InstaNovo-P identified 86.75% more peptides that contained phosphorylated sites compared to the labeled space (Figure 5A). The distribution of these peptides matched well with the standard peptide distribution of human peptides in MS data^24^ and the database search space (Figure 5B). We also explored InstaNovo-P predictions in the labeled and full search space in relation to the database search. At 5% FDR, we predicted more than double the peptides compared to the model predictions that overlap with the database search in the same threshold (Figure 5C).

**Figure 5.**
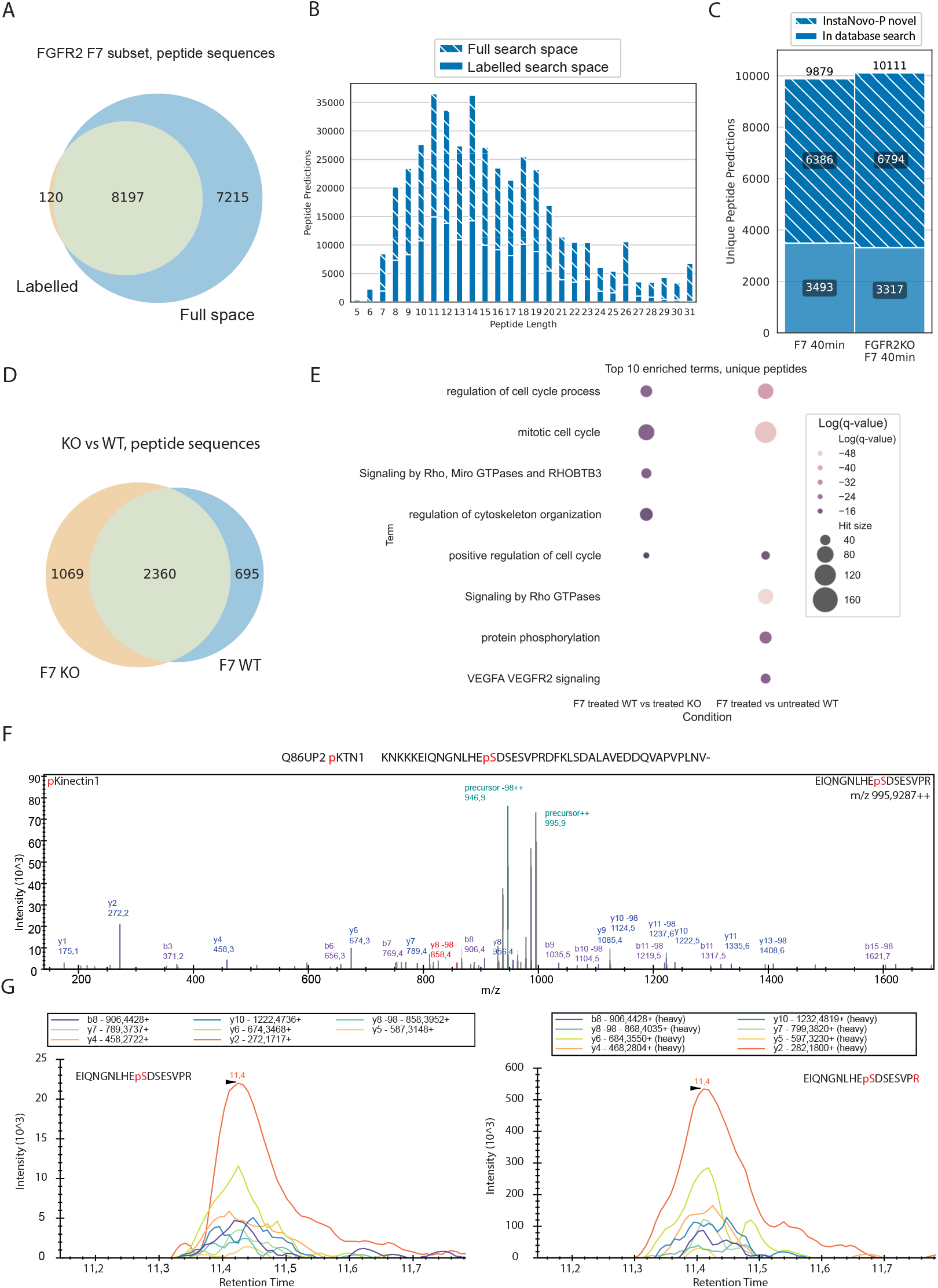
InstaNovo-P predicts biologically relevant phosphorylation events. A) Venn diagram of the number of phosphorylated peptides in the database search space (labeled), and predicted phosphorylated peptides from InstaNovo-P in the FGFR2 dataset, F7 treated subset at 5% FDR. B) Histogram of unique phosphorylated peptide predictions at 5% FDR as a function of peptide length. C) Peptides predicted by InstaNovo-P 5% FDR, labeled by presence or absence in database search. D) Unique peptides predicted by InstaNovo-P at 5% FDR in WT and KO replicates, when treated with F7 for 40 minutes. E) Selected enriched gene ontology terms in the two conditions (F7 treated WT vs KO, and WT treated vs untreated, at 40 minutes) of the FGFR2 dataset. F) Best scan for the phosphorylated peptide “EIQNGNLHEpSDSESVPR” in the WT cells. G) Fragment ion transitions for the same precursor peptide for endogenous (light peptide, left) and reference peptide (heavy peptide, right)

When comparing the predictions between F7-treated KO and WT cells at the 40 min, InstaNovo-P identified 695 peptides that were unique for WT (Figure 5D) at 5% FDR. These peptides may represent the core of F7-FGFR2 signaling in our experimental model as they were found in the WT but not in the KO cells. Out of these, 153 were not found in the database search space, with 138 of them found only in this condition and timepoint. Importantly, 10 out of 10 of these novel top predicted phosphorylated sites, when ranked by number of PSMs, represented known phosphorylated sites in PhosphoSitePlus^10^. As an example, we identified a phosphorylation site at S402, in the C-terminal peptide of SAC31, a protein predicted to be involved in centrosome duplication and mitotic progression (the most commonly phosphorylated residue in this protein according to experiments integrated in PhosphoSitePlus). This suggests that our model is able to identify known and validated phosphorylated peptides, providing complementary information and substantially improving phosphoproteomics searches.

The comparison between F7-treated vs untreated WT cells revealed 2,340 phosphorylated peptides unique for the treated samples, with 417 identified only by InstaNovo-P, and 368 of these identified only in that condition. Similarly to the WT versus KO comparison, all 10 top ranked novel peptides contained phosphorylated sites present in PhosphoSitePlus. Three of these 10 peptides mapped to nucleophosmin (NPM1), and contain the same phosphorylated site at position S125, possibly phosphorylated by the kinases Aurora A or B. This site is the most frequent phosphorylation event in NPM1, and mutations in this position show mitotic defects^38^. This phosphorylation event causes the protein to shift to a monomeric state instead of a pentameric^39^, and re-localize from the nucleolus to midbody^38^. However, as none of these peptides identified on NPM1 are tryptic, they would not have been identified with the standard database search settings. These peptides differed by one residue at the N-terminus (starting with GSGPVHISGQHIVAVEEDAES(+79.97)EDEEEEDVK), with several cleavage events downstream of this peptide, hinting at aminopeptidase degradation of the protein in a sequential way. Quantification of these peptides corroborated this observation, with the truncated peptidoforms found in increasing or diminishing abundance (Supplementary Figure S5). Although caspases -3, -7 and -8 can cleave NPM1 during apoptosis^40^, no cleavage or aminopeptidase activity has been reported for that protein region. We also observed an unmodified, cleaved product in the acidic region of NPM1 (161-188) with cleavage at position 180. This suggests that InstaNovo-P can detect modified, non-tryptic peptides and provide another layer of information in proteomics searches.

We further scrutinized the novel predictions in these two experimental conditions, by performing gene enrichment analysis for the proteins that the novel phosphorylated peptides map to with Metascape^41^ and default settings with human background. This analysis revealed expected processes significantly enriched in these conditions such as regulation of the cell cycle and RHO GTPase activity (Figure 5E). Overall, these results indicate that our model successfully predicted phosphorylated peptides that recapitulate the known biological processes downstream of F7-FGFR2 signaling in T47D cells^42–44^.

To confirm our observations, we validated the predicted phosphorylated serine residue on kinectin-1 (KTN1) at position 75 with targeted proteomics in newly collected samples for the same conditions. We designed a parallel reaction monitoring (PRM) assay for the phosphorylated peptide “EIQNGNLHEpSDSESVPR” within residue 66 to 82 we identified with our predictions but not with database search. We detected the peptide precursor with the 98 dalton mass shift on the y8 ion which indicated the loss of the phosphoric acid group through fragmentation, confirming our *de novo* predictions (Figure 5F). Furthermore, we could show that the ion chromatograms of the reference peptide (isotopically labeled with C13 arginine at the C-terminus) and the retention time aligned well with the endogenous peptide within WT T47D breast cancer cells (Figure 5G).

These results show that applying InstaNovo-P to FGFR2 phosphoproteomics uncovered hundreds of additional phosphorylation sites, many of which map to known signaling proteins and are catalogued in phosphorylation databases. Enrichment analysis tied these discoveries to cell-cycle events, and targeted proteomics confirmed the potential of InstaNovo-P in predicting biologically meaningful phosphorylation events that traditional database searches miss.

## Discussion

In this work, we introduce and evaluate the phosphorylation-specific *de novo* peptide sequencing model, InstaNovo-P. We find InstaNovo-P to increase performance of the base model, InstaNovo, in unmodified peptides in the AC-PT test set and 21PTM-unmod dataset. This indicates that catastrophic forgetting during fine-tuning was mitigated as the model not only maintains, but improves performance in the source domain. We believe that the encoder-first gradual unfreezing schedule contributed to this effect. By unfreezing the encoder layers before the decoder, we allowed the model to adapt to phosphorylation-specific features while preserving generalisable representations from the base model. This strategy may have helped avoid catastrophic forgetting, as reflected in improved recall on both phosphorylated and unmodified peptides. Further, our model was not only successful at improving detection of phosphorylated peptides, but also improving general sequencing performance. We speculate that this performance gain stems from InstaNovo not having achieved ceiling performance and benefitting from further training. It is also possible that seeing examples of modified peptides, or data from diverse origins, improves learning of unmodified peptide representations.

InstaNovo-P delivered state-of-the-art recall on phosphorylated peptide PSMs. InstaNovo-P was able to detect novel peptides with high confidence, thus discovering phosphorylated peptides not detected by database search. Importantly, these phosphorylation events have been detected in previous proteomics experiments, validating our approach and findings. However, database search still identifies a substantial number of peptides not detected with InstaNovo-P, at least not as the top prediction. This is potentially due to lack of a complete fragment ion series, low fragment ion signal, chimeric spectra, or single residue errors in the peptide predictions. Overall, we believe that InstaNovo-P can be used in tandem with database searches to increase phosphorylated peptide identification.

We also find a substantial number of InstaNovo-P predictions that map to proteins in the reference proteome while not present in database results, further validating our predictions. Additionally, its confidence scores correlate with true FDR and database localization certainty, offering a powerful complementary layer of site-localization information. We observed several instances of our model disagreeing with database localization assignment, and validating these cases will be part of future work. We believe that tighter integration with database search engines, where InstaNovo-P could rescue ambiguous or unassigned spectra, will maximize phosphoproteome coverage and accelerate biological discovery.

InstaNovo-P shows diminished performance in peptides with more than one phosphorylation sites. Similarly, we see a correlation between peptide recall and peptide length. Longer sequences tend to result in increased sequence complexity and ambiguity, as well as potential missed cleavages, which make them harder to identify. Additionally, longer peptides often suffer from lower ionization efficiency, making them more difficult to detect in mass spectrometry. Unsurprisingly, we observe best performance on phosphorylation events in serine residues and worst performance on tyrosine phosphorylation events, reflecting our training data composition. This bias, where performance drops for peptides with more phosphorylation sites and less abundant phosphorylation events, can be explained by the increased complexity of the task and imbalance of data. We expect that our performance in threonine and tyrosine phosphorylation events will catch up once the model has been trained with adequate data for these residues.

InstaNovo-P exhibits its highest recall on the 21PTM test set, potentially because it is synthetic peptides closely resembling our training dataset for the base model. Admittedly, it is challenging to compare models trained on different datasets with potential for data leakage. In this work, we have implemented a strict homology based split to control data leakage.

InstaNovo-P has fixed inference, in contrast to database search were the space increases with addition of modifications. This might be beneficial for computational costs and search times, especially when large reference proteomics and additional modifications are considered. InstaNovo-P is only compatible with data dependent acquisition, as it relies on additional information about precursor mass and charge. Adapting the model, or preprocessing data independent acquisition spectra to be compatible with the model will be part of future directions. Similarly, we expect part of future work to be the interpretation of InstaNovo-P weights and attention, that can possibly inform on fragmentation events and ions that enable the model to accurately predict phosphorylated peptides.

Finally, in this study we describe a reproducible framework providing a blueprint for model fine-tuning for different modifications or application domains with substantial biological utility, thus facilitating broader advancements within proteomics research.

## Methods

### Data preprocessing

For a description of the All-Confidence ProteomeTools (AC-PT) dataset used to train the base model, see the InstaNovo^24^ paper. The dataset used for fine tuning is comprised of a collection of reprocessed PRIDE projects in Scop3P^31^ (For a list of the projects, see Supplementary Table 1). The dataset originally contains 4, 053, 346 PSMs. To only fine-tune on high confidence PSMs, the dataset is filtered at a confidence threshold of 0.80, reducing it to 2, 760, 939 PSMs, representing 74, 686 unique peptide sequences. Most of the data is of human origin, except for PXD005366 and PXD000218, which contain a mix of human and mouse. All PSMs that were used to train the model contained at least one phosphorylated site, while 169, 114 PSMs (6%) contained oxidated methionine. The entries were acquired using a mixture of instruments, whereas the AC-PT dataset was acquired using only Orbitrap Fusion.

To partition the fine-tuning dataset into training, validation and test, GraphPart^32^, an algorithm for homology partitioning, was applied on the set of unique peptide sequences. GraphPart was set to use MMseqs2^45^ with a partitioning threshold of 0.8 and a train-validation-test ratio of 0.7*/*0.1*/*0.2. Of the 74, 686 unique sequences, 390 were removed by GraphPart, reducing the total number of PSMs to 2, 691, 117 in a 2, 008, 923*/*232, 641*/*449, 553-split, although during training, a random subset of only 2% of the validation set was used in order to reduce computation.

The 21PTM^34^ dataset consists of around 5000 synthetic peptides carrying 21 different post-translational modifications as well as their unmodified counterparts. We used the phosphorylated and unmodified subsets. To minimize data leakage, we removed any sequences that overlapped with the fine-tuning training set. We removed sequences that had identical sequences, even if they had different phosphorylation sites. For the phosphorylation subset, we dropped 4094 sequences, going from 45252 to 41158. For the unmodified subset, we dropped 2188 sequences, going from 34823 to 32635. Even though we removed sequences that overlapped with our phosphorylation training-set, 21PTM still contains a significant overlap with the training set of AC-PT, used to train the base InstaNovo model. Specifically, 56.15% of unmodified sequences in 21PTM overlap with AC-PT, owing to the fact that 21PTM is part of the ProteomeTools project. Of course, none of the phosphorylated sequences overlap. For evaluation of predictions in all datasets, we remapped carbamidomethylated cysteines (C[+57.02]) to “C”, and isoleucine (I) to leucine (L).

### Fine-tuning

Before fine-tuning, we reinitialized the head and sequence embedding of the model decoder, as they had to contain the extended vocabulary of the 3 phosphorylation tokens. All parameters in the specific layers were randomly initialized, therefore not carrying over any information from previous learning.

To fine-tune the base InstaNovo^24^ model we implemented a novel encoder-first gradual unfreezing fine tuning schedule. This gradual unfreezing^33^ approach is typically implemented by first freezing the entire model, and then unfreezing layers, starting at the head, then progressing from the top of the decoder to the bottom of the decoder and sequence embedding, then from top to bottom through the encoder. We differ from this approach by first unfreezing the head and sequence embedding, then gradually the encoder, then gradually the decoder. We also explored other methods of fine-tuning, including full-model whereby the entire model is unfrozen from the start, head-only where only the final classifier layer is unfrozen and head-then-full where once the head has converged, the remainder of the model is unfrozen. When applying head-only fine-tuning, we also had to reinitialise the sequence embedding layer due to the different vocabulary sizes of the fine-tuning dataset and the dataset the base model was trained on. The alternative fine-tuning approach of encoder-first gradual unfreezing was motivated by the fact that the more standard fine-tuning approaches quickly led to overfitting.

We find the Slanted Triangular Learning Rate (STLR)^33^ scheduler to be most performant for this encoder-first gradual unfreezing. The learning rate is warmed up for the first 10% of the STLR period, and annealed for the remainder of the period (See Supplementary Figure S1A). We applied the STLR scheduler in a cyclical manner, whereby each phase of unfreezing restarts the STLR schedule, thus allowing for all frozen parameters to warmup. This warmup strategy was found to be most performant (see Supplementary Table 4 for hyperparameters and Supplementary data for the learning-rate schedule used). The best model was found in the final epoch at step 273,725 at a validation loss of 0.3949.

The primary Python packages used for fine-tuning were PyTorch^46^, Lightning^47^, and Fine-Tuning Scheduler^48^ to manage the gradual unfreezing and learning rate scheduling.

### Metrics

We evaluated our model using peptide-level metrics: peptide precision, peptide recall and Area-Under-Curve (AUC) of precision-recall curves and recall-coverage curves. We used exact string matching to decide whether two peptides were equal. For further details, see the InstaNovo^24^ paper.

### Cell culture and FGFR2 depletion

The human epithelial cell line T47D (HTB-133) was purchased from ATCC and authenticated shortly before the samples were collected through short tandem repeat (STR) analysis of 21 markers by Eurofins Genomics, yielding positive results. The cells were routinely monitored monthly for mycoplasma contamination using a PCR-based detection assay (Venor® GeM–Cambio). T47D cells were cultured in RPMI 1640 Medium, GlutaMAX™ Supplement (Gibco), supplemented with 2 mM L-glutamine, 100 U/ml penicillin, 100 *µ*g/ml streptomycin, and 10% fetal bovine serum. For gene editing, guide RNAs (crRNA) specific to FGFR2 (FGFR2 crRNA 1: GCCCTACCTCAAGGTTCTCA; FGFR2 crRNA 2: ACCTTGAGAACCTTGAGGTA) were synthesized (IDT) and combined with a common trans-activating crRNA (Alt-R® CRISPR-Cas9 tracrRNA, IDT) to form a functional ribonucleoprotein (RNP) duplex. The resulting complexes were pre-incubated with Cas9 nuclease (Alt-R® S.p. Cas9 Nuclease V3, IDT) and transiently transfected into parental T47D cells using Viromer® CRISPR transfection reagent (Cambridge Bioscience), following the manufacturer’s protocol. Colonies were subsequently selected and screened for genomic editing efficiency using the Inference of CRISPR Edits (ICE) analysis tool provided by Synthego. Successful knockout of FGFR2 protein expression was validated by Western blot analysis^19^. T47D were serum starved overnight and treated with 100 ng/mL of FGF7 or FGF10 from Peprotech for 40 min or 72h before lysis for MS analysis.

### Proteomics sample preparation

T47D samples for proteomics and phosphoproteomics were prepared as follows^19^: Cells were washed with PBS and lysed in ice-cold 1% Triton lysis buffer supplemented with Pierce protease inhibitor tablets (Life Technologies) and phosphatase inhibitors (5 nM Na3VO4, 5 mM NaF, and 5 mM *β* -glycerophosphate). Proteins were precipitated overnight at ™ 20 ^*°*^C in a four-fold excess of ice-cold acetone. Acetone-precipitated proteins were then solubilized in a denaturation buffer consisting of 6 M urea and 2 M thiourea in 10 mM HEPES (pH 8). Cysteines were reduced with 1 mM dithiothreitol (DTT) and alkylated with 5.5 mM chloroacetamide (CAA). Proteins were subsequently digested using endoproteinase Lys-C (Wako, Osaka, Japan) followed by sequencing-grade modified trypsin (Sigma), and the digestion reaction was quenched using 1% trifluoroacetic acid (TFA). Peptides were purified using reversed-phase Sep-Pak C18 cartridges (Waters, Milford, MA) and eluted with 50% ACN.

A small fraction (1%) of the eluted peptides was reserved for proteome analysis. These peptides were evaporated using a speed vacuum, resuspended in 40 *µ*L of 0.1% TFA, 5% ACN, and loaded onto C18 STAGE-tips. For phosphoproteomic analysis, 6 mL of 12% TFA in ACN was added to the remaining peptides, which were then enriched using TiO_2_ beads (5 *µ*m, GL Sciences Inc., Tokyo, Japan). The beads were suspended in 20 mg/mL 2,5-dihydroxybenzoic acid (DHB), 80% ACN, and 6% TFA, and the samples were incubated in a 1:2 (w/w) sample-to-bead ratio for 15 min with rotation. After centrifugation for 5 min, the supernatant was collected and subjected to a second incubation with a two-fold dilution of the previous bead suspension. Elution of phosphorylated peptides was done with 20 *µ*L of 5% NH_3_, followed by 20 *µ*L of 10% NH_3_ in 25% ACN, which were evaporated to a final volume of 5 *µ*L in a speed vacuum. The concentrated phosphorylated peptides were acidified by adding 20 *µ*L of 0.1% TFA and 5% ACN, and loaded onto C18 STAGE-tips. Peptides were eluted from STAGE-tips in 20 *µ*L of 40% ACN, followed by 10 *µ*L each of 60% ACN and 100% ACN, then reduced to 5 *µ*L by SpeedVac. Finally, 5 *µ*L of 0.1% FA and 5% ACN was added. A 1 *µ*L aliquot of the sample (for proteome analysis) or a 3 *µ*L aliquot (for phosphoproteome analysis) was transferred to a 5 *µ*L sample loop and loaded onto the column at a flow rate of 300 nL/min with 5% B for 5 and 13 minutes, respectively. After sample loading, the loop was removed from the flow path, and the flow rate was decreased from 300 nL/min to 200 nL/min within 1 minute, with the mobile phase adjusted to 7% B.

### Mass spectrometry data acquisition and analysis

Purified peptides were analyzed by LC-MS/MS using an UltiMate® 3000 Rapid Separation LC system (RSLC, Dionex Corporation, Sunnyvale, CA) coupled to a Q Exactive HF mass spectrometer (Thermo Fisher Scientific, Waltham, MA). Mobile phase A consisted of 0.1% formic acid (FA) in water, and mobile phase B contained 0.1% FA in acetonitrile (ACN). Chromatographic separation was performed using a 75 mm × 250 *µ*m inner diameter, 1.7 mM CSH C18 analytical column (Waters). Peptides were separated using a linear gradient as follows: 7–18% B over 64 minutes, 18–27% B over 8 minutes, and 27–60% B in 1 minute. The column was subsequently washed at 60% B for 3 minutes and re-equilibrated for 6.5 minutes. At 85 minutes, the flow rate was increased back to 300 nL/min until the end of the run at 90 minutes. Mass spectrometry data were acquired over 90 minutes in positive ion mode using data-dependent acquisition. Peptides were automatically selected for fragmentation using a Top8 method (phosphoproteome) or Top12 method (proteome), based on precursors with charge states of 2, 3, or 4 and an m/z range of 300–1750 Th. Dynamic exclusion was set to 15 s. MS1 resolution was 120,000, with an AGC target of 3 *×* 10^6^ and a maximum fill time of 20 ms. For MS2, resolution was set to 60,000 with an AGC target of 2 *×* 10^5^ and a maximum fill time of 110 ms (Top12), or 30,000 with the same AGC target and a 45 ms fill time (Top8). The isolation window was set to 1.3 Th, and normalized collision energy was 28.

Raw data were analyzed using the MaxQuant software suite (https://www.maxquant.org; version 1.6.2.6) with the integrated Andromeda search engine. Proteins were identified by searching the HCD-MS/MS peak lists against a target/decoy version of the human UniProt Knowledgebase database, including complete proteome sets and isoforms (version 2019; https://uniprot.org/proteomes/UP000005640_9606), supplemented with commonly observed contaminants such as porcine trypsin and bovine serum proteins. Tandem mass spectra were matched with a mass tolerance of 7 ppm for precursor ions and 0.02 Da (or 20 ppm) for fragment ions. Carbamidomethylation of cysteine was set as a fixed modification. Variable modifications included protein N-terminal acetylation, N-pyro-glutamine formation, oxidized methionine, and phosphorylation of serine, threonine, and tyrosine for phosphoproteome analyses. For proteome datasets, variable modifications included N-terminal acetylation, oxidized methionine, and deamidation of asparagine and glutamine. Site localization probabilities were computed by MaxQuant using the PTM scoring algorithm. Data were filtered by posterior error probability to ensure an FDR below 1% at the levels of peptides, proteins, and modification sites. Only peptides with an Andromeda score greater than 40 were included in the final analysis.

### Quantification of InstaNovo-P predictions

Quantification was performed using AUC calculation for the predictions at their measured mass charge (m/z), with a tolerance of ±0.05. PSMs with identical sequences, but a difference in mass of more than ±0.05, were considered different PSMs. For identical PSMs, the retention time (RT) and m/z from the first identified PSM was used for quantification. Boundaries for the AUC were defined to limit cross-interference as the largest window between local minima within a ±1% RT window of the identification RT. Quantification was performed on MS1 spectra. Noise was calculated for each experiment, as the mean of the lowest 5% intensities from all MS2 spectra in the given file, and then subtracted from each intensity measurement during calculation of AUC. PSMs were grouped across experiment files if mass was within tolerance of ±0.05 and sequences and modifications were identical. Each group was then considered as one peptide. Peptide mass was determined by the average mass of the combined PSMs, and confidence was assigned as the highest confidence of the peptide’s PSMs.

### Targeted proteomics validation

Confluent (80%) T47D breast cancer cells were starved in FBS-free medium for 24 hours. Subsequently cells were either treated with 100 ng/mL FGF7 or kept untreated for 40 minutes. Cells were placed on ice and 100 µL of lysis buffer (4 M GuHCl and 100 mM HEPES (pH 8.0)) were added together with Pierce protease inhibitor tablets (Life Technologies) and cells were scraped off. Scraped cells were transferred into a fresh tube, heated to 95^*°*^C for 5 minutes, and sonicated at 30% for 30 seconds while kept on ice. Lysed cells were centrifuged at 13,000 x g for 15 minutes at 4^*°*^C and the supernatant was transferred into a fresh tube. The Bradford assay was used to determine protein concentration and 5 *µ*g of protein lysate were used for reduction with Tris-(2-carboxyethyl)-phosphin (TCEP) (5 mM final concentration) and alkylation with CAA both carried out at 400 rpm, 65^*°*^C for 30 minutes. After reduction and alkylation, samples were diluted 8-fold to reduce the GuHCl concentration to 0.5 M using 100 mM HEPES (pH 8.0) and Lys-C (Wako, Osaka, Japan) (1:100) was added for 4 hours followed by overnight digestion with sequencing-grade modified trypsin (Sigma) at 37^*°*^C. The reaction was quenched using 1% TFA. Peptides were purified using Solaµ plates (Thermo Fisher Scientific, Waltham, MA). Purified peptides and phosphorylated peptides were analyzed by LC-MS/MS using an Vanquish LC system (Thermo Fisher Scientific, Waltham, MA) coupled to a Exploris 480 mass spectrometer (Thermo Fisher Scientific, Waltham, MA). Mobile phase A consisted of 0.1% formic acid (FA) in water, and mobile phase B contained 0.1% FA in 80% acetonitrile (ACN). Chromatographic separation was performed using a µPAC HPLC analytical column (Thermo Fisher Scientific, Waltham, MA). Peptides were separated using a gradient as follows: 1-10% B over 4.1 minutes, 10.1–22.5% B over 19 minutes, 22.5-45% over 8.4 minutes, and 45-95% over 6.5 minutes. The flow rate is kept at 750 nL/min over the initial 4.1 minutes and changed to 250 nL/min over 33.9 minutes. An initial data-dependent acquisition (DDA) strategy is used to acquire the spectral library of selected peptides and different charge states. The mass list for the reference heavy peptides is DINNIDYYK (583,284402 m/z), DINNIDYYK (389,192027 m/z), DINNIDYYK (292,145839 m/z), DINNIDY[+80]Y[+80]K (663,250733 m/z), DINNIDY[+80]Y[+80]K (442,502914 m/z), DINNIDY[+80]Y[+80]K (332,129005 m/z), EIQNGNLHESDSESVPR (960,949709 m/z), EIQNGNLHESDSESVPR (640,968898 m/z), EIQNGNL-HESDSESVPR (480,978493 m/z), EIQNGNLHES[+80]DSESVPR (1000,932875 m/z), EIQNGNLHES[+80]DSESVPR (667,624342 m/z), EIQNGNLHES[+80]DSESVPR (500,970075 m/z), GISSLPR (370,220455 m/z), GISSLPR (247,149395 m/z), GISSLPR (185,613865 m/z), GISS[+80]LPR (410,20362 m/z), GISS[+80]LPR (273,804839 m/z), GISS[+80]LPR (205,605448 m/z). The scans were unscheduled and a dependent scan was performed on the most intense ion if no target was found. For MS1, resolution was set to 120000 with an AGC target of 300% or 3×10e6 and a maximum fill time of 50 ms over a scan range of 300-1500 m/z. For MS2 resolution was set to 7500, isolation window was set to 1 m/z, normalized collision energy was set to 30% over a scan range of 150-1700 m/z and the AGC target was 1000% or 1×10e6 and a maximum fill time of 10 ms.

Following the DDA scan the peptides were acquired using a unscheduled parallel reaction monitoring (PRM) strategy within the biological matrix to obtain retention times, using the following mass list for endogenous and reference heavy peptides: DINNIDY[+80]Y[+80]K (659.243634 m/z), DINNIDY[+80]Y[+80]K[+8] (663.250733 m/z), DINNIDYYK (579.277303 m/z), DINNIDYYK[+8] (583.284402 m/z), EIQNGNLHES[+80]DSESVPR (995.92874 m/z), EIQNGNL-HES[+80]DSESVPR[+10] (1000.932875 m/z), EIQNGNLHESDSESVPR (955.945575 m/z), EIQNGNLHESDSESVPR[+10] (960.949709 m/z), GISSLPR (365.21632 m/z), GISSLPR[+10] (370.220455 m/z), GISS[+80]LPR (405.199486 m/z), and GISS[+80]LPR[+10] (410.20362 m/z). For MS1, resolution was set to 120000 with an AGC target of 300% or 3×10e6 and a maximum fill time of 246 ms over a scan range of 300-1500 m/z. For MS2 resolution was set to 7500, isolation window was set to 1 m/z, normalized collision energy was set to 27% over a scan range of 150-1700 m/z and the AGC target was 1000% or 1×10e6 and a maximum fill time was set to auto.

Raw data were analyzed via Skyline^49^ (Version 24.1.0.414, Seattle, WA). The DDA survey run was imported and converted into a spectral library. The spectral library was built using the directed DDA runs, including following modifications, Carbamidomethyl (C), phosphorylation (STY) and isotope modifcations at the C-terminus (13C(6)15N(4) (C-term R) and 13C(6)15N(2) (C-term K)). Light and heavy peptides were included and unscheduled runs were used to detect retention times that were exported and used for scheduled PRM runs. MS1 filtering was performed by including a minimum of 20% of the base peak at 10 ppm accuracy. MS2 filtering was set to PRM as acquisition method.

## Data Availability

The InstaNovo base model has been trained on the ProteomeTools datasets, part I-III and can be found in the ProteomeXchange Consortium via the PRIDE partner repository^29^ with identifiers PXD004732 (Part I), PXD010595 (Part II) and PXD021013 (Part III). The 21-PTM dataset (ProteomeTools part V) can also be found in PRIDE under the identifier PXD009449. The FGFR2 dataset has been deposited in PRIDE under dataset identifier PXD062859. Reviewers can access the FGFR2 dataset with username and password `eZorng4dZASn.` The targeted proteomics experiment files have been deposited to PanoramaWeb and can be accessed with the link https://panoramaweb.org/4W3hr8.url with reviewer and password `4^+NDURy%8H!Al.`

The InstaNovo-P training dataset can be found on HuggingFace at https://huggingface.co/datasets/InstaDeepAI/InstaNovo-P. Supplementary data supporting the data preprocessing and analysis performed on our training and validation datasets can be found on figshare at 10.6084/m9.figshare.29049533.

## Code Availability

Inference code and model checkpoints for the base InstaNovo model are available at the InstaNovo GitHub repository https://github.com/instadeepai/instanovo. The checkpoint for InstaNovo-P v1.0.0 is available as an [asset to InstaNovo release v1.1.2](https://github.com/instadeepai/InstaNovo/releases/tag/1.1.2) as well as a Jupyter notebook describing the usage of InstaNovo-P. Code used for fine-tuning will be shared upon publication. Custom scripts used for data analysis and visualization are available upon request and will also be uploaded to a public repository upon publication.

## Acknowledgements

K.K. is supported by a Novo Nordisk Foundation Young Investigator Award (grant no. NNF16OC0020670) and a postdoctoral fellowship grant from the Independent Research Fund Denmark (grant no. 4257-00010B). K.K. and E.M.S. also acknowledge support from PRO-MS: Danish National Mass Spectrometry Platform for Functional Proteomics (grant no. 5072-00007B). C.F., J.F., and P.F. are supported by Wellcome Trust (107636/ Z/15/Z and 107636/Z/15/A). C.F. is also supported by a Novo Nordisk Foundation Young Investigator Award (grant no. NNF22OC0070845). V.C. is supported by the LEO Foundation (LF-OC-23-001220 to C.F.). We are grateful to the DTU Proteomics Core Facility for maintenance and running of instruments. We especially thank M. Wennekers Nielsen and M. Vestegaard Lukassen for their help and advice during troubleshooting, sample preparation and data acquisition.

## Author contributions statement

K.K. and T.P.J. conceived the project. P.R., T.C, S.v.P, L.M. and C.F. provided datasets for training and validation. J.L. preprocessed the data, trained the model, and performed benchmarking with feedback from R.C., A.M., K.E., J.V.G, E.M.S., T.P.J and K.K. P.F. and J.F. performed cell culture, FGFR2 depletion, sample preparation and mass spectrometry, and C.F. searched the proteomics raw data. J.L. used the trained model to perform de novo sequencing in the validation datasets. A.K.M. and I.S.G. performed quantification of the FGFR2 dataset de novo predictions. V.C., C.F. and K.K. analysed the FGFR2 data. K.K. and J.L. performed proteome mapping and downstream analysis. N.L.C., J.V.G., T.P.J. and K.K. supervised the project. J.L., P.R., R.C., T.P.J. and K.K. drafted the original manuscript. All authors reviewed the manuscript.

## Supplementary Tables

**Table 1.** PRIDE Accession codes used for training, validation and test sets

|  |
| --- |
| PXD006482<sup>50</sup> |
| PXD005366<sup>51</sup> |
| PXD004447<sup>52</sup> |
| PXD004940<sup>53</sup> |
| PXD004452<sup>54</sup> |
| PXD004415<sup>55</sup> |
| PXD004252<sup>56</sup> |
| PXD003198<sup>57</sup> |
| PXD003657<sup>58</sup> |
| PXD003531<sup>59</sup> |
| PXD003215<sup>60</sup> |
| PXD002394<sup>61</sup> |
| PXD002286<sup>62</sup> |
| PXD002255<sup>63</sup> |
| PXD002057<sup>64</sup> |
| PXD001565<sup>65</sup> |
| PXD001550<sup>66</sup> |
| PXD001546<sup>66</sup> |
| PXD001060<sup>67</sup> |
| PXD001374<sup>68</sup> |
| PXD001333<sup>69</sup> |
| PXD000474<sup>70</sup> |
| PXD000612<sup>71</sup> |
| PXD001170<sup>72</sup> |
| PXD000964<sup>73</sup> |
| PXD000836<sup>74</sup> |
| PXD000674 |
| PXD000680<sup>75</sup> |
| PXD000218<sup>76</sup> |

**Table 2.**
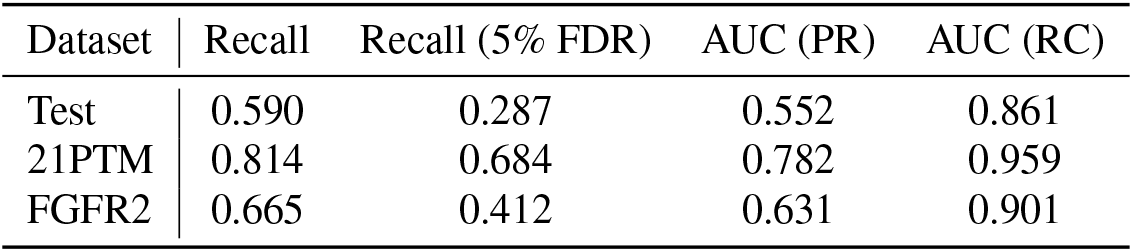
InstaNovo-P zero-shot evaluation on 3 datasets

| Dataset | Recall | Recall (5% FDR) | AUC (PR) | AUC (RC) |
| --- | --- | --- | --- | --- |
| Test | 0.590 | 0.287 | 0.552 | 0.861 |
| 21PTM | 0.814 | 0.684 | 0.782 | 0.959 |
| FGFR2 | 0.665 | 0.412 | 0.631 | 0.901 |

**Table 3.** Peptide recall- and precision for peptides containing phosphorylation at different amino acids

| Dataset | S Recall | S Precision | T Recall | T Precision | Y Recall | Y Precision |
| --- | --- | --- | --- | --- | --- | --- |
| Test | 0.603 | 0.644 | 0.493 | 0.545 | 0.421 | 0.473 |
| 21PTM | 0.000 | 0.000 | 0.000 | 0.000 | 0.812 | 0.824 |
| FGFR2 | 0.688 | 0.712 | 0.539 | 0.566 | 0.417 | 0.453 |

**Table 4.** Hyperparameters

| Hyperparameter | Value |
| --- | --- |
| Learning Rate<br>(per epoch) | [2e-4, 1e-5, 5e-6, 5e-6, 5e-6,<br>5e-6, 2e-6, 2e-6, 2e-6, 2e-6] |
| STLR ratio | 0.03125 |
| STLR cycle length | 29591 |
| STLR midpoint fraction | 0.1 |
| Weight decay | 1e-6 |
| Dropout | 0.1 |
| Train Batch Size | 64 |
| Epochs | 10 |
| Gradient Accumulation | 1 |
| Gradient Clip Value | 10.0 |
| Train Subset | 1.0 |
| Valid Subset | 0.02 |
| Predict Batch Size | 2220 |
| Validation Check Interval | 7397 |
| Number of Peaks | 200 |
| Min M/Z | 50.0 |
| Max M/Z | 2500.0 |
| Min Intensity | 0.01 |
| Remove Precursor Tolerance | 2.0 |
| Max Charge | 10 |
| Precursor Mass Tolerance | 50 ppm |
| Isotope Error Range | [0, 1] |
| Model Dimension | 768 |
| Number of Heads | 16 |
| Feedforward Dimension | 1024 |
| Number of Layers | 9 |
| Number of Beams | 2 |

## Supplementary Figures

**Figure S1.**
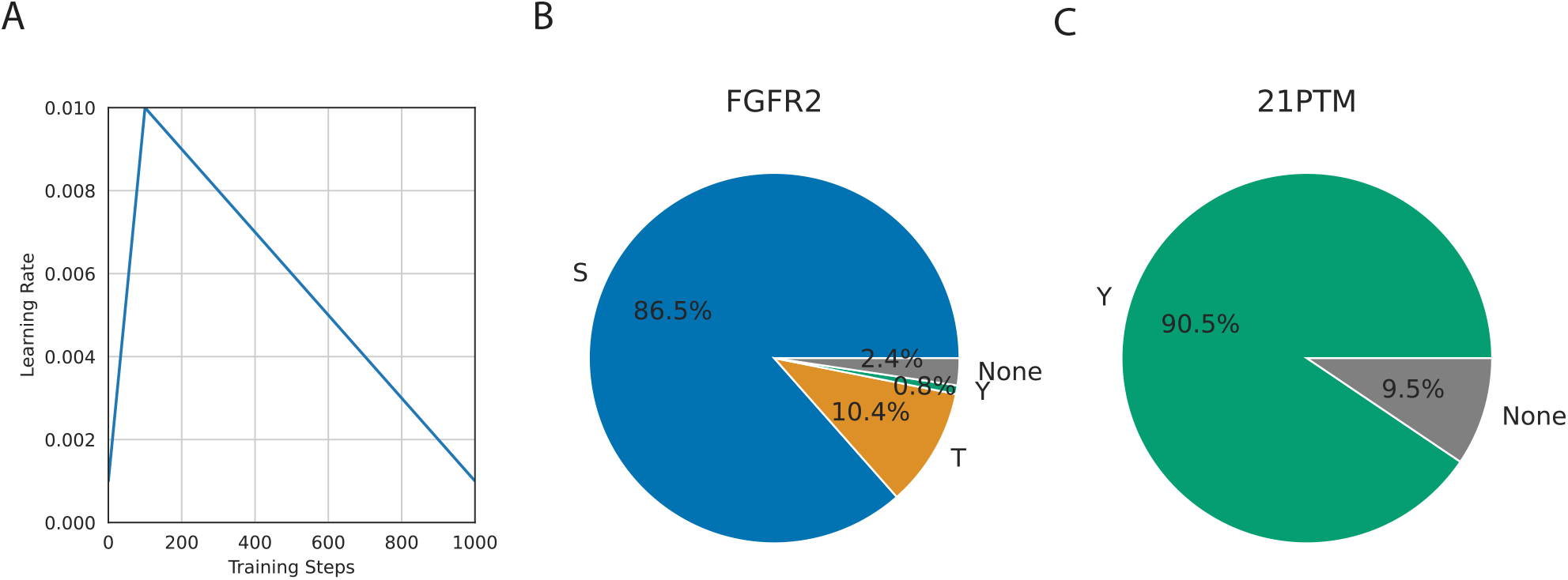
Supplementary figure 1. A) Learning rate during base model finetuning and InstaNovo-P training. B) Proportion of phosphorylation events per residue type in the FGFR2 and C) the 21PTM dataset.

**Figure S2.**
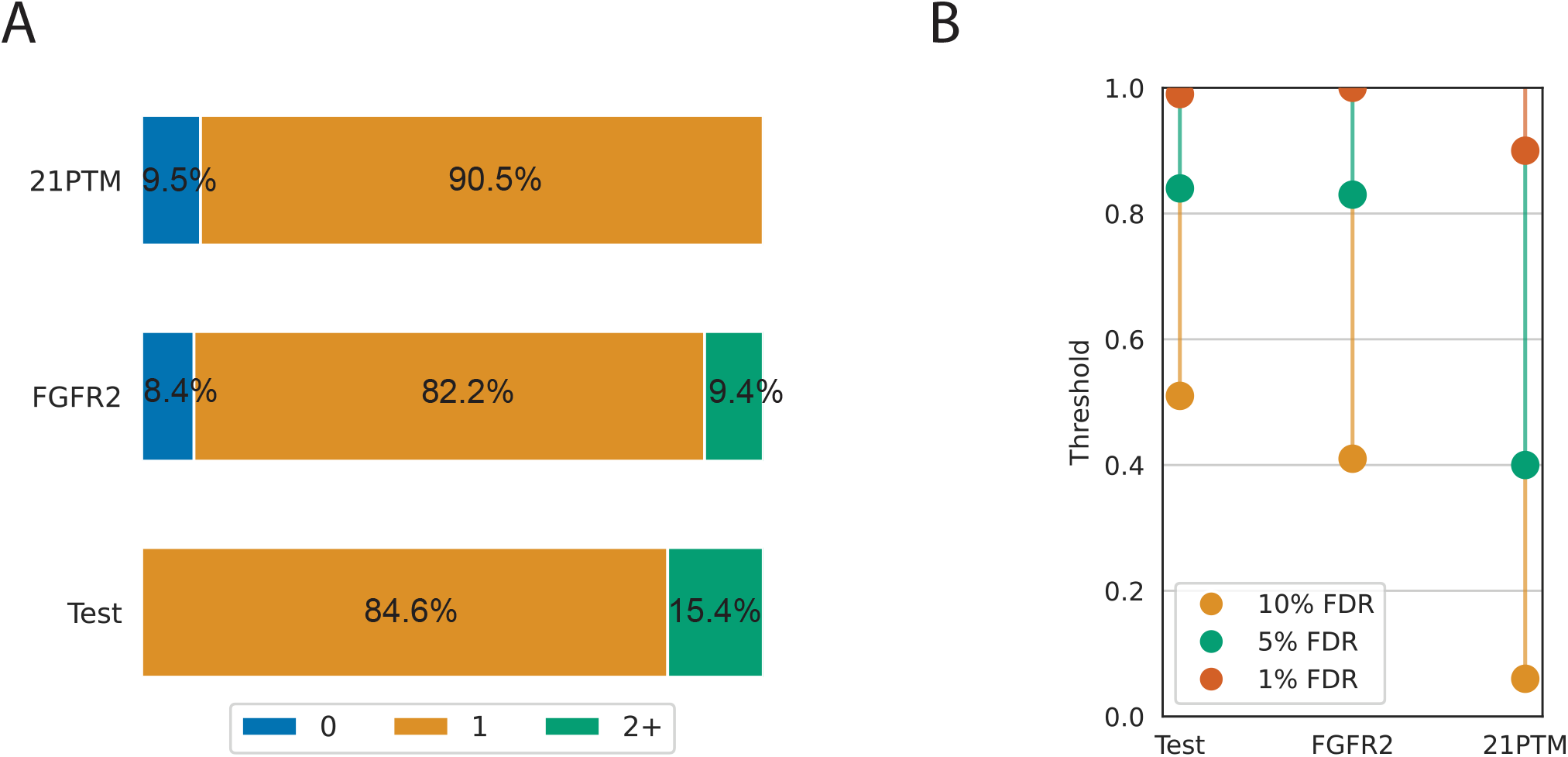
Supplementary figure 2. A) Proportion of PSMs with 0 (blue), 1 (orange), and 2 phosphorylated sites (green) in the 3 test datasets used. B) Computed confidence thresholds for different FDR in our 3 datasets.

**Figure S3.**
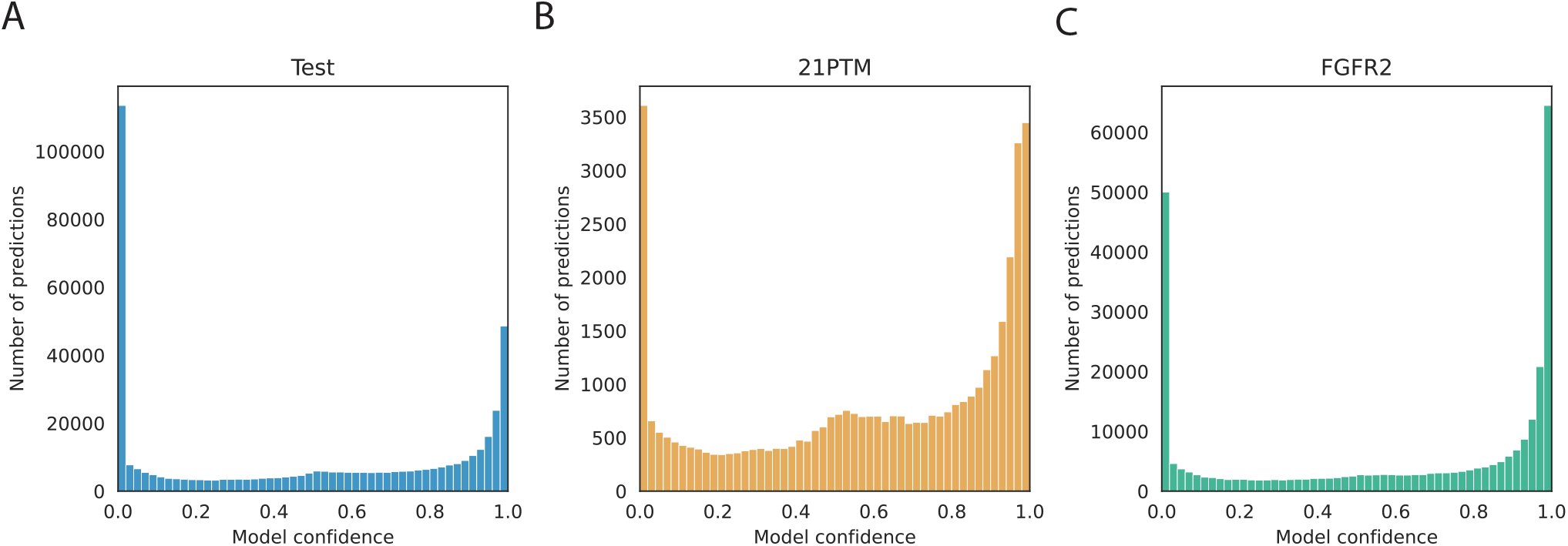
Supplementary figure 3. A) Confidence distribution for peptide predictions of InstaNovo-P in our test dataset, B) the 21PTM dataset, and C) the FGFR2 dataset.

**Figure S4.**
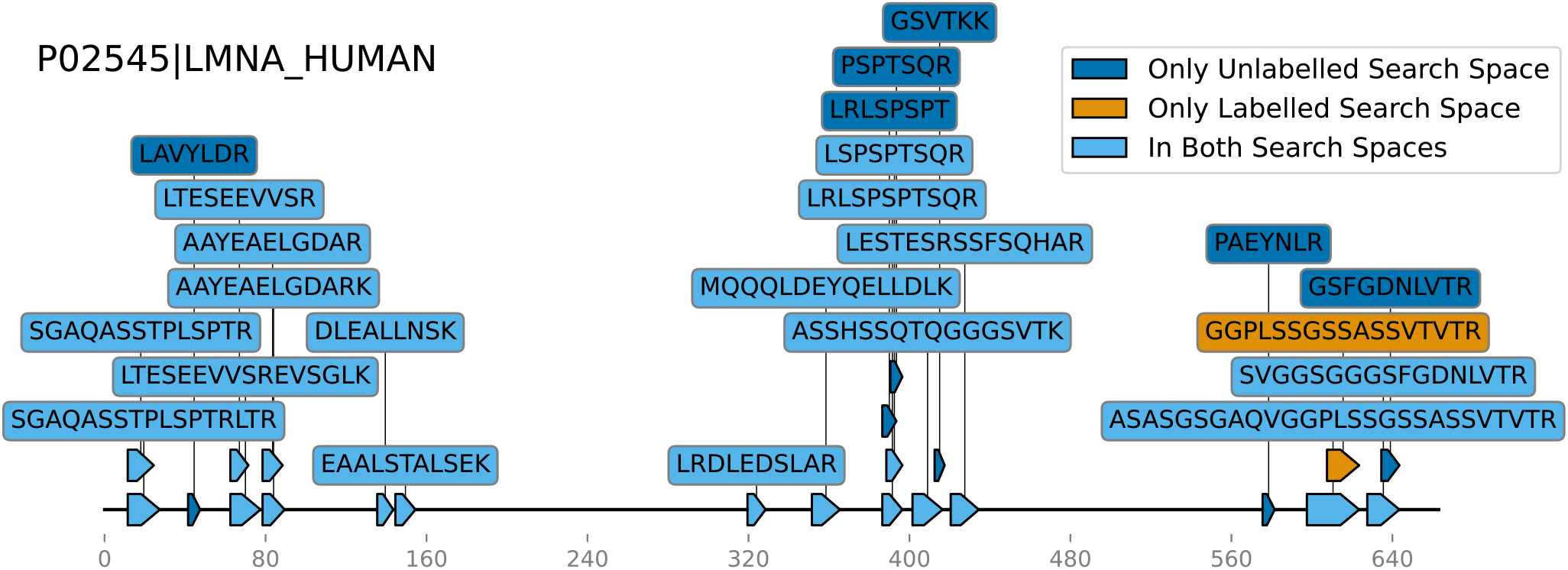
Supplementary figure 4. Mapping of phosphorylated peptides identified with database search (bottom), and novel phosphorylated peptides identified with InstaNovo-P in LMNA (Uniprot accession P02545) at 5% FDR.

**Figure S5.**
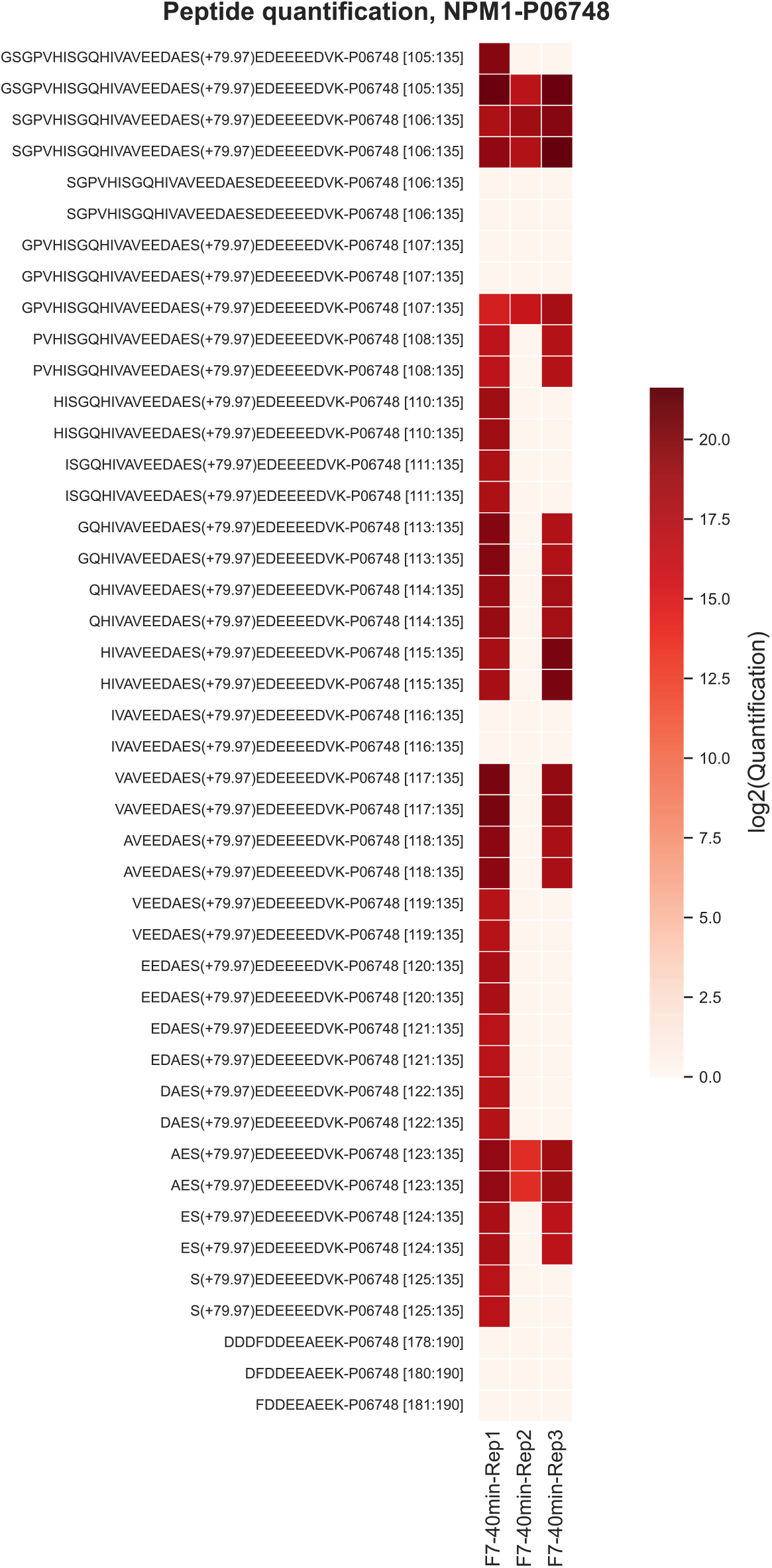
Supplementary figure 5. Quantification of semi-tryptic phosphorylated and non-phosphorylated peptides detected with InstaNovo-P in protein NPM1 (Uniprot accession P06748) at 5% FDR.

